# A video-rate hyperspectral camera for monitoring plant health and biodiversity

**DOI:** 10.1101/2024.01.18.576183

**Authors:** Laura J. Brooks, Daniel Pearce, Kenton Kwok, Nikhil Jawade, Man Qi, Erola Fenollosa, Deniz Beker, James Whicker, Katrina Davis, Roberto Salguero-Gómez, Robin Wang, Steve Chappell

## Abstract

Hyperspectral cameras are a key enabling technology in precision agriculture, biodiversity monitoring, and ecological research. Consequently, these applications are fuelling a growing demand for devices that are suited to widespread deployment in such environments. Current hyperspectral cameras, however, require significant investment in post-processing, and rarely allow for live-capture assessments. Here, we introduce a novel hyperspectral camera that combines live spectral data and high-resolution imagery. This camera is suitable for integration with robotics and automated monitoring systems. We explore the utility of this camera for applications including chlorophyll detection and live display of spectral indices relating to plant health. We discuss the performance of this novel technology and associated hyperspectral analysis methods to support an ecological study of grassland habitats at Wytham Woods, UK.

## 1. INTRODUCTION

Hyperspectral imaging technologies are highly valuable for applications in the plant sciences, ecology, and agriculture. Indeed, plant tissues exhibit strong signatures in the visible and near-infrared regions of the electromagnetic spectrum, which correlate with plant health, growth, and productivity.^1^ Analyses of these spectral signals can therefore be used to infer important information about a plant’s condition, such as leaf chlorophyll content,^2–6^ nutrient status or deficiency,^7–9^ tissue damage,^10^ and the presence of stressors such as drought, disease, and pests.^11–17^

In the agricultural sector, spectral data can enable ‘precision farming’.^18^ In precision farming, data provided by sensing platforms (e.g., unmanned aerial vehicles [UAVs], autonomous robots, planes, satellites) is used to inform decision-making and management practices, as well as the automation of key processes such as harvesting, yield estimation, and fertiliser application.^16^, ^19^ In this context, hyperspectral imaging has the potential to facilitate high-throughput crop monitoring and quality control, as well as informing the judicious and efficient use of water, chemical fertilisers, pesticides and other interventions.^20, 21^ The widespread deployment of affordable hyperspectral imaging devices in agriculture therefore promises numerous benefits, including reduced costs for producers, increased food security, and mitigation of the harmful environmental effects associated with some conventional farming practices.^22^

In ecology, hyperspectral sensing can be utilised to measure and monitor vegetation health,^2, 23^ and to study the complex effects of environmental factors and human activities on biodiversity.^24–26^ Faced with global climate change and other pressing environmental concerns, such as increased pollution and soil degradation, the ability to gather accurate information about the effects of such factors on biodiversity in natural settings is becoming ever more critical. There is, therefore, a strong motivation to explore the potential of hyperspectral tools and techniques for ecology research and environmental monitoring. Particularly, the field is currently lacking suitable devices that provide high resolution data at low-cost, that are accessible, easy to use, and can be validated *in situ*. Progress in this area is expected to leverage recent advancements in machine learning and computer vision to enable the automated analysis of large hyperspectral datasets,^20^ thereby placing a high demand on the compute power required of any such hyperspectral imaging and analytical platform.

In Section 2 of this paper, we introduce in the Living Optics camera: a novel, video-rate, snapshot hyperspectral imager. This new technology has the potential to address current bottlenecks to the application of hyperspectral imaging in plant science, biodiversity monitoring, and ecological research: affordability, portability, live assessment, ease of use, and integration with advanced computational analysis techniques. The utility of this camera to infer information about plant health is then explored in Section 3, with a focus on the detection of chlorophyll and the mapping of relevant spectral indices. Finally, in Section 4, we present our early work on deploying the camera, and associated analysis methods, to support the *in situ* study of grassland habitats at Wytham Woods, an iconic ecological monitoring site managed by the University of Oxford, UK. The paper concludes with a discussion of future work and the outlook for this emerging hyperspectral device.

## 2. VIDEO-RATE HYPERSPECTRAL IMAGING

Much of the prior work on spectral imaging for agriculture and environmental monitoring has utilised airborne or space-based spectral systems. These classical approaches have traditionally focused on multispectral or line-scanned hyperspectral imaging.^18, 27^ Multispectral line-scanned imaging systems typically incorporate a series of wavelength-dependent filters aligned to specific pixels on a 2D imaging array, with the array being scanned across the scene.^28, 29^ Line-scanned hyperspectral imaging systems typically consist of an imaging slit, positioned at the focal plane of an objective lens, relay optics, and a dispersive element in front of a 2D array of detector pixels. In this traditional format, the slit images the scene along one axis across the detector, while the dispersive element disperses the wavelengths across the other axis. To build up a hyperspectral image from this approach, the slit must then be scanned across the scene.^29, 30^ Scanning is typically achieved using the inherent motion of the platform on which the camera is mounted (e.g. UAV, satellite, etc). It is possible to translate line-scanning techniques from airborne/space-based viewing geometries to ground-based or laboratory environments by incorporating additional mechanisms to scan the scene, such as a mirror or scanning table. However, scanning limits the rate that sequential hyperspectral data cubes can be captured, and line-scan techniques are generally only suited to static scenes with well-characterised camera motion and fixed illumination conditions.

An alternative to line-scanning systems are ‘snapshot’ spectral imagers.^29, 31^ Snapshot systems typically trade spatial resolution, spectral resolution, or noise performance to achieve instantaneous hyperspectral data cubes without scanning. The collection modes of scanning and snapshot hyperspectral systems are compared diagrammatically in Figure 1. It is expected that the practical benefits and video-rate analytics afforded by snapshot hyperspectral imaging will outweigh the tradeoffs *versus* line-scanning hyperspectral imaging, for a wide range of applications. Although video-rate images are not traditionally required for plant science applications (e.g. Lowe *et al*, 2017^11^), due to the relatively static nature of the subject matter, the instantaneous user feedback afforded by snapshot techniques provides unique opportunities for data collection efficiency and *in situ* targeted scientific discovery, in addition to improving the practicalities of pointing, focusing, and capturing imagery in the field.

**Figure 1.**
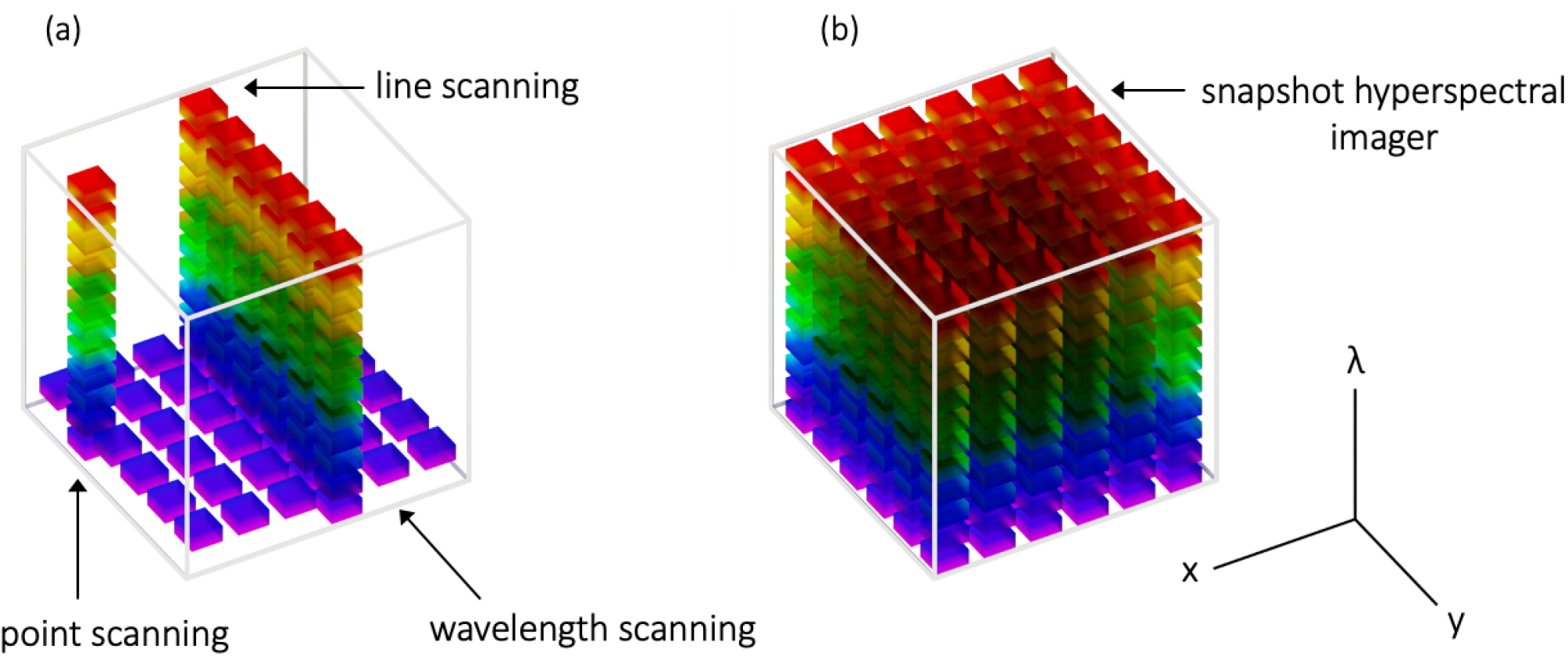
A comparison of the hyperspectral data collection modes for (a) scanning hyperspectral imagers, and (b) snapshot hyperspectral cameras. The diagrams indicate the data captured during a single integration time, for each type of imaging system. For the scanning approaches shown in panel (a), multiple exposures are required, in order to build up a full hyperspectral dataset. By comparison, snapshot hyperspectral cameras (b) collect a full hyperspectral data cube in a single exposure.

**Figure 2.**
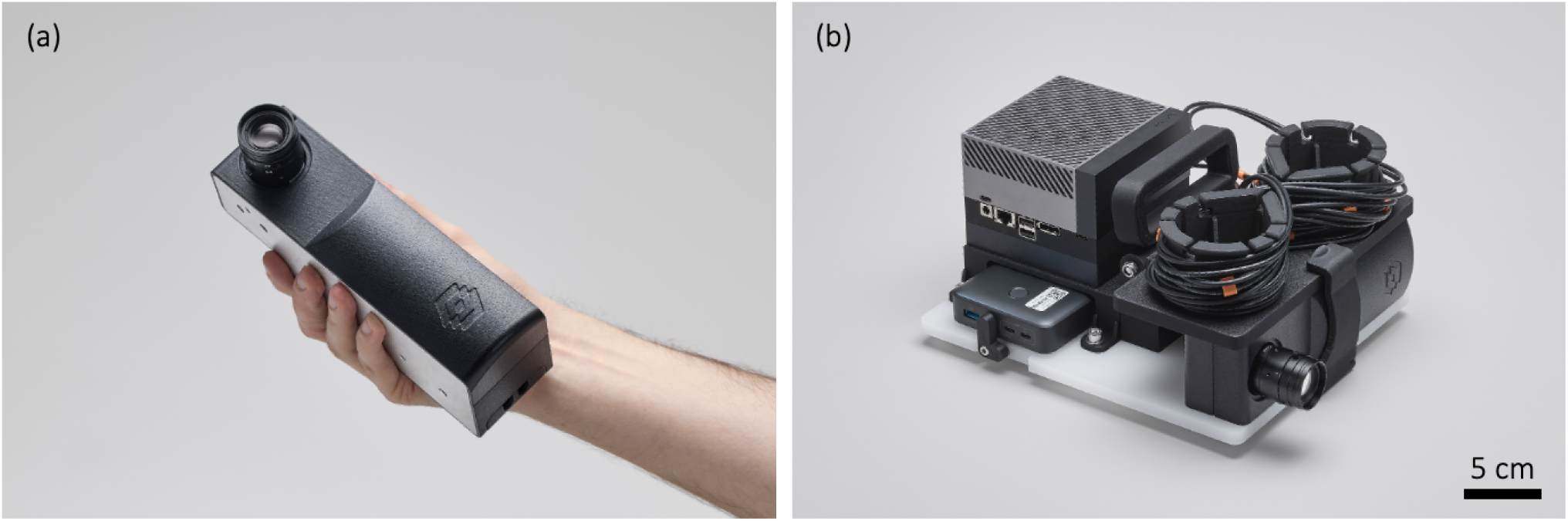
Photographs showing the Living Optics hyperspectral camera. (a) The camera module, which is sufficiently compact as to be held in one hand. (b) The camera development kit, which includes a camera module, edge compute (Jetson AGX Orin, NVIDIA), power bank, lenses, cabling and portable chassis.

The newly-available video-rate snapshot hyperspectral camera presented here enables handheld, close-range hyperspectral applications, bringing these traditionally expensive and sophisticated tools to a wider audience, and facilitating applications in numerous fields. The Living Optics camera produces two, axially-aligned outputs from the hyperspectral imaging system - a 2048 x 2432 pixel ‘scene view’ image, and a set of 4384 spectra, evenly sampled from across the scene. The spectra from each of the 4384 sampling points contain 96 bands, dispersed between 440 nm and 900 nm. To enable live applications on these two data streams, arriving at up to 30 Hz, as well as instantaneous feedback to the users, an NVIDIA Jetson AGX Orin has been paired with the camera. All of the applications discussed in this paper have been captured and processed using this combined edge compute and camera development kit, fitted with a 35 mm lens with an approximate minimum focal distance of 30 cm.

## 3. PLANT HEALTH ANALYSIS

In this section, we describe the application of the Living Optics camera for analysing plant health, using both conventional spectral indices and custom metrics devised from our own hyperspectral analysis.

### 3.1 Mapping spectral indices pertaining to plant health

A common tool in the interpretation of hyperspectral data is the use of spectral indices.^11^ A spectral index is a formula that takes as input selected wavebands from the spectral information, and returns a single value that is designed to assess a quantity of interest. Applying a spectral index calculation to the spectra at each (x,y) spatial coordinate in a 3-dimensional (x, y, channels) hyperspectral dataset allows the data to be summarised in a two-dimensional array that can be conveniently displayed as an image, referred to here as a ‘spectral index map’. This approach is widespread in remote sensing,^32^ and can also be applied to close-range, or ‘proximal’ hyperspectral measurements.^33^

A great number of spectral indices pertaining to plant health have been explored in the literature, with the aim of quantifying a range of factors such as vegetation density and health,^34^ leaf chlorophyll content,^3, 5^ carotenoid concentration,^4, 6^ and many others.^2, 12, 13, 19, 35, 36^ In Figure 3, we demonstrate the use of the Living Optics camera to obtain and display such spectral indices. The example spectral index maps show a scene containing indoor plants, analysed using five well-established indices relating to plant health. These indices are: (1) Normalised Difference Vegetation Index (NDVI);^34^ (2) Modified Chlorophyll Absorption in Reflectance Index (MCARI);^3^ (3) Red-edge Chlorophyll Index (RECI);^5^ (4) Photochemical Reflectance Index (PRI);^35, 36^ and (5) Carotenoid Concentration Index (CRI700).^4, 6^ The formulae for each of these indices, together with the wavebands applied to compute them using Living Optics data, are presented in Table 1. Each index is calculated using reflectance information from the scene, therefore, a reflectance conversion step is required prior to spectral index calculation. This calculation transforms the raw radiance data captured by the camera into a reflectance measurement, *R*(*λ*), normalised between 0 and 1, where 0 means that no light is reflected at this wavelength, and 1 means that all of the light of this wavelength incident on the object is reflected. To perform this conversion, for each wavelength *λ*, the radiance values, *S*(*λ*), collected from the scene are divided by the intensity of the illuminant, *I*(*λ*), falling on the objects in the scene, according to the equation:

**Table 1.** Details of the spectral indices used to generate the plots shown in Figure 3. The ‘Wavebands’ column lists the wavelength limits between which the reflectance values were extracted and averaged, to obtain the inputs for the spectral index calculation.

| Index | Index name | Formula | Wavebands<br>(start, stop) | Citation |
| --- | --- | --- | --- | --- |
| NDVI | Normalised Difference Vegetation Index | $\frac{R_{NIR} - R_{RED}}{R_{NIR} + R_{RED}}$ | $R_{RED}:(665,675);$<br>$R_{NIR}:(795,805);$ | Rouse (1974)<sup>34</sup> |
| MCARI | Modified Chlorophyll Absorption in Reflectance Index | $[(R_{700} - R_{670}) - 0.2 \times (R_{700} - R_{550})] \times \frac{R_{700}}{R_{670}}$ | $R_{550}:(545,555);$<br>$R_{670}:(665,675);$<br>$R_{700}:(695,705)$ | Daughtry (2000)<sup>3</sup> |
| RECI | Red-edge Chlorophyll Index | $\frac{R_{750}}{R_{710}} - 1$ | $R_{750}:(745,755);$<br>$R_{710}:(705,715)$ | Gitelson (2005)<sup>5</sup> |
| PRI | Photochemical Reflectance Index | $\frac{R_{531} - R_{570}}{R_{531} + R_{570}}$ | $R_{531}:(526,536);$<br>$R_{570}:(565,575)$ | Gamon (1992, 1997)<sup>35, 36</sup> |
| CRI700 | Carotenoid Concentration Index | $\frac{1}{R_{515}} - \frac{1}{R_{700}}$ | $R_{515}:(510,520);$<br>$R_{700}:(695,705)$ | Gitelson (2003a, 2006)<sup>4, 6</sup> |

**Figure 3.**
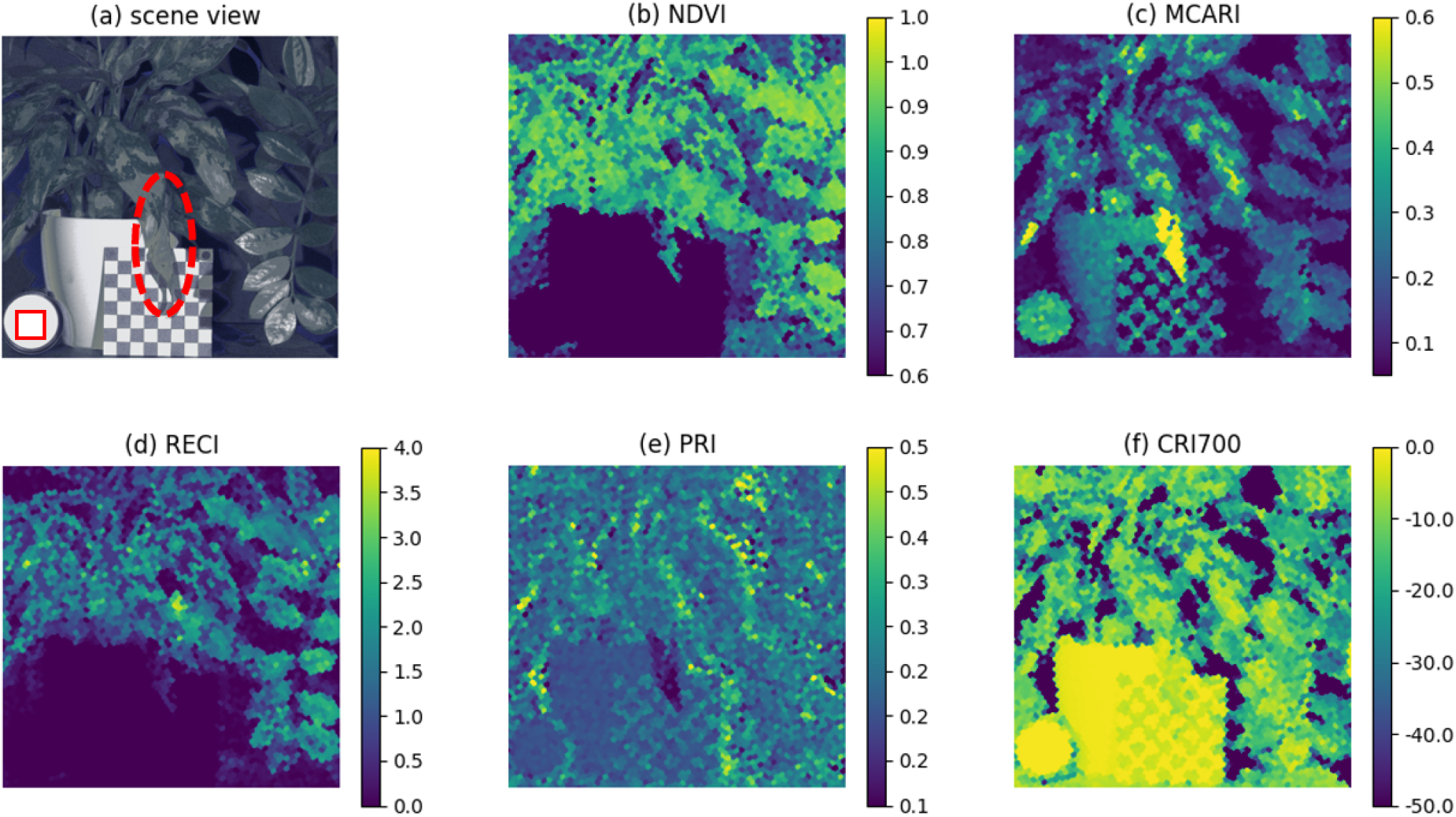
Various spectral indices plotted for a single scene containing two indoor plants, *Aglaonema ‘Maria’* (left of image) and *Zamioculcas zamiifolia* (right of image). The RGB ‘scene view’ image is presented in panel (a). A solid red square in the ‘scene view’ image indicates the locations from which the reference spectra were extracted for reflectance conversion. Panels (b-f) show maps plotted for the following spectral indices: (b) NDVI - Normalised Difference Vegetation Index, (c) MCARI - Modified Chlorophyll in Reflectance Index, (d) RECI - Red-edge Chlorophyll Index, (e) PRI-Photochemical Reflectance Index, and (f) Carotenoid Concentration Index. Definitions of the indices used to generate the spectral index maps are given in Table 1. The colour scale limits in these maps have been selected to optimise the contrast in parts of the scene containing plant matter. For scale, each square of the the checkerboard included in the image is 12.7 x 12.7 mm.

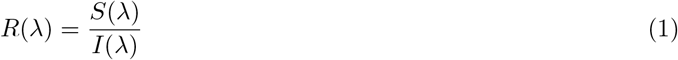

To evaluate the illuminant spectrum, it is necessary to include a uniformly reflecting white object in the image. In the example shown in Figure 3, a standard white reflectance tile (LabSphere, Spectralon diffuse reflectance standard, white, 99%) is placed in the scene for this purpose. A reference spectrum *I*(*λ*) is then obtained by taking the mean of all the spectra extracted from the locations within the solid red square indicated on the scene view image in the figure.

As shown in Figure 3, each map highlights different features of the scene according to the index plotted. We note that there is strong contrast, in each of the spectral maps, between the apparently healthy plant leaves, and the wilted, ‘unhealthy’ leaf, indicated by the dashed oval in the scene view panel. Given that all of the plotted indices are expected to correlate with plant health, this spectral index contrast gives a positive indication of the potential of the Living Optics camera for this application. In the following sections, we further explore the application of the camera for plant health assessment, undertaking quantitative measurements to investigate the camera’s utility for quantifying leaf chlorophyll content, a key indicator of plant health.

### 3.2 Quantification of chlorophyll content in solution

To investigate the use of the Living Optics camera to detect chlorophyll, we first performed measurements on solutions containing known concentrations of chlorophyll extract in water. By varying the concentration of these solutions and analysing the resulting spectral response, a custom metric was proposed to indicate the chlorophyll content. A calibration curve was then plotted, linking this metric to the concentration of the chlorophyll extract in the solution.

Samples were prepared by diluting commercially-obtained chlorophyll extract (Bioactive Liquid Chlorophyll, Four-Leaf Farmacy, UK) in water. Six samples ranging from 1.3 - 9.1 % chlorophyll extract by volume were studied. We note that these percentages refer to the dilution of the extract in water, rather than an absolute measure of the chlorophyll concentration. A 100% water sample was used as a reference for normalisation of the spectra, and a sample of green food dye was also measured for comparison.

Transmission illumination geometry was chosen for this experiment to allow for comparison with standard techniques such as transmission spectrophotometry.^4^ Broadband, even illumination, was achieved using a white light source, fibre-coupled to a small integrating sphere, as shown in Figure 4. Cuvettes, with 10 mm path length, containing the solutions to be measured, were placed in front of the opening of the integrating sphere. Hyperspectral images were then obtained using a Living Optics camera fitted with a 35 mm focal length objective lens. During data collection, the setup was covered with a black cloth to prevent any reflection of ambient light from the samples.

**Figure 4.**
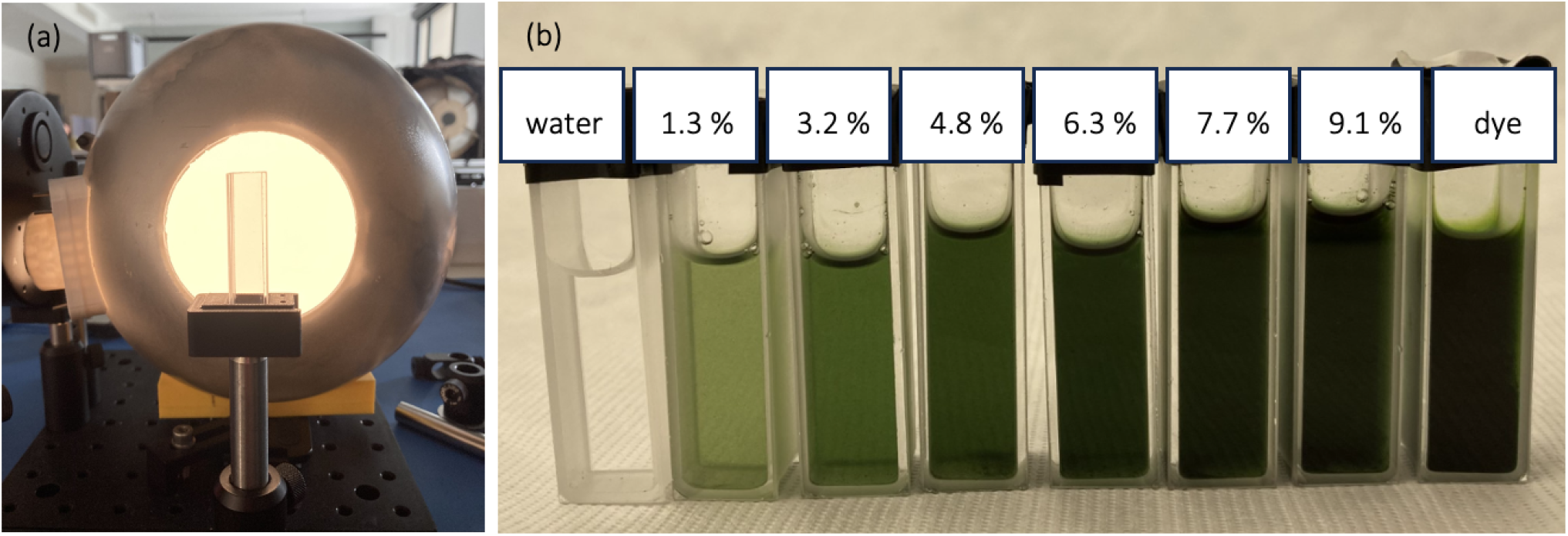
Investigating detection of chlorophyll in solution via the Living Optics hyperspectral camera. (a) Experimental set up for transmission imaging. A broadband fibre light source (not pictured) illuminates a small integrating sphere as shown. Cuvettes are placed sequentially in front of the opening of the sphere and imaged using a Living Optics hyperspectral camera. (b) Samples measured. Cuvettes containing different concentrations of chlorophyll extract diluted in water. For comparison, a sample of green dye was also measured.

Transmission spectra, plotted in Figure 5(a), were extracted from the hyperspectral images of each sample. As shown in the figure, the chlorophyll solutions are clearly distinguishable from the green food dye according to their spectral shape. For all chlorophyll samples, a strong absorption (indicated by a dip in the transmission spectrum) is observed in the orange/red region of the spectrum, around 640 nm. This result was confirmed independently, using a point spectrometer measurement of the same solutions.

**Figure 5.**
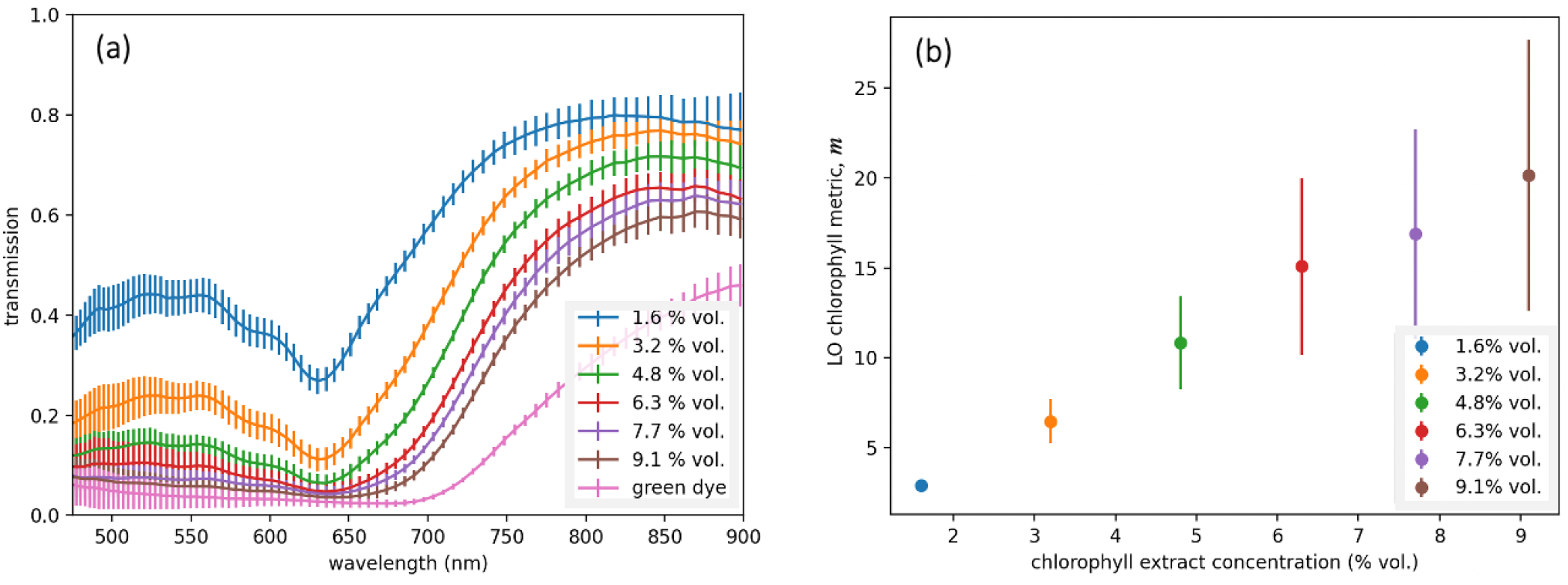
Quantifying chlorophyll content in solution using hyperspectral data (a) Transmission spectra for samples of different chlorophyll extract concentrations. Each spectrum is the mean over approx. 60 spectra extracted within the region of interest in the hyperspectral image. Spectra are normalised to the transmission of a 100 % water sample. (b) Calibration curve showing the mean value of the proposed metric, *m*, calculated for each of the extracted spectra, plotted against extract concentration. In both (a) and (b), the error bars indicate *±* one standard deviation of the datasets used to calculate the mean.

To quantify the chlorophyll content, a custom chlorophyll metric, *m*, was devised. This metric computes the ratio between T_610*−*660_, the average transmission in the 610 – 660 nm range (i.e., the region of spectrum where the chlorophyll samples are observed to be strongly absorbing), and T_850*−*900_, the average transmission in a reference region in the near-infrared (850 – 900 nm):

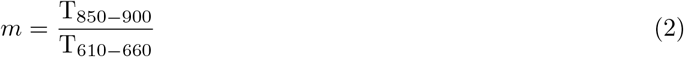

As shown in Figure 5(b), this metric shows good correspondence with the concentration of the chlorophyll samples, within the range of concentrations tested. This suggests that such an approach could therefore be used as an indicator of chlorophyll level for practical applications using Living Optics hyperspectral data.

However, further investigation and optimisation of the metric may improve these results. In particular, it is noted that the relationship between the metric and the concentration does not show the logarithmic behaviour expected from the Beer-Lambert Law. This is thought to be related to background subtraction methods and is a matter for future investigation. Nevertheless, these results provide confidence that hyperspectral data obtained by the camera can be used to assess chlorophyll concentration, and on this basis we proceed in Section 3.3 to explore the camera’s utility to quantify leaf chlorophyll content.

### 3.3 Quantification of leaf chlorophyll

To investigate the detection of chlorophyll within plant leaves, four *Butterhead* lettuce leaf samples were prepared as shown in Figure 6. To ensure maximum variation in leaf chlorophyll content across the samples, four leaves were selected from different parts of the plant, ranging from the outermost to the innermost leaves. A total of nine regions of interest (ROIs) were selected from across the four leaf samples, to include the greenest regions at the tips of the leaves, through to the palest regions close to the base and stem. The ROIs, with dimensions approximately 18 x 18 mm, were marked onto the leaves using black tape, and their locations are indicated by the black rectangles shown in Figure 6.

**Figure 6.**
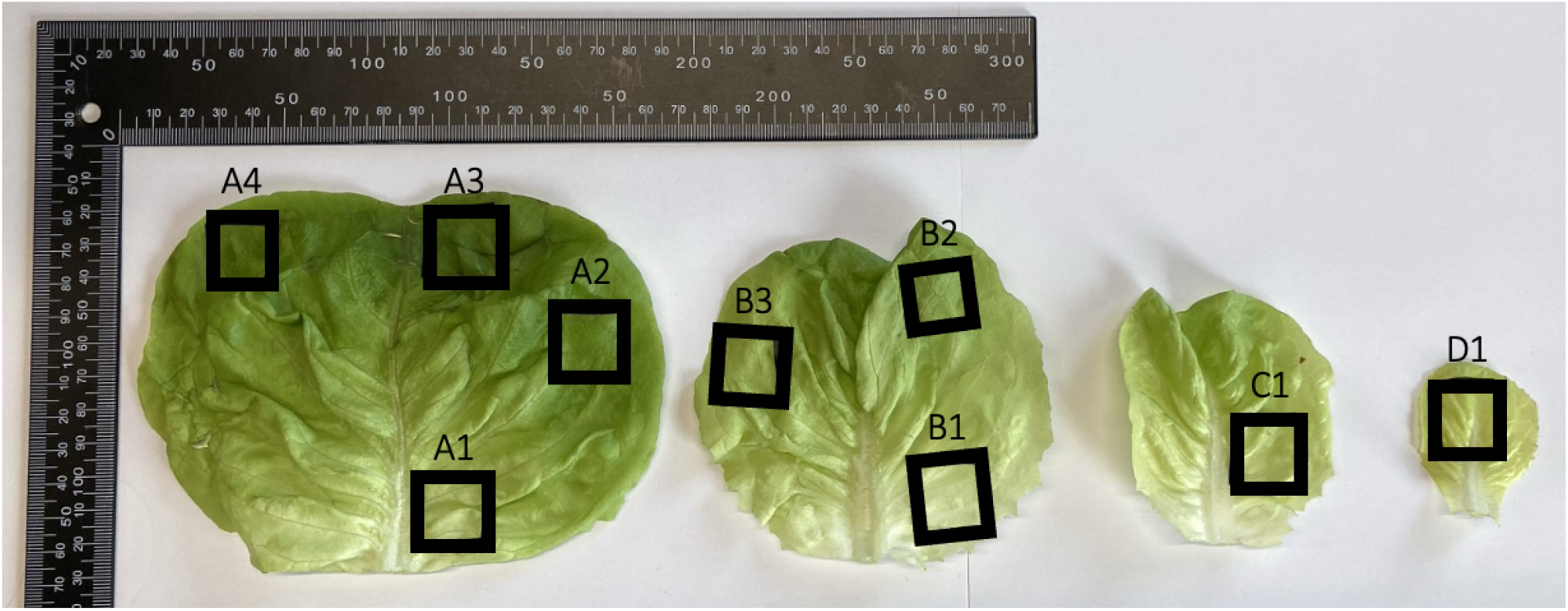
Photograph showing lettuce leaf samples measured for chlorophyll analysis, using both the Living Optics camera, and a commercially-obtained chlorophyll ‘SPAD’ meter. Black rectangles indicate the locations of the regions of interest analysed.

The leaf samples were imaged using a Living Optics hyperspectral camera, and a broadband halogen light source. A standard reflectance tile (LabSphere, Spectralon diffuse reflectance standards, white, 99%) was also imaged under the same lighting conditions and exposure settings, to enable characterisation of the illumination for the purpose of computing a reflectance conversion, which was performed in post processing. In the analysis of the hyperspectral data, approximately 200 spectra were extracted from within each leaf ROI. The resulting mean reflectance spectrum for each region is plotted in Figure 7(a).

**Figure 7.**
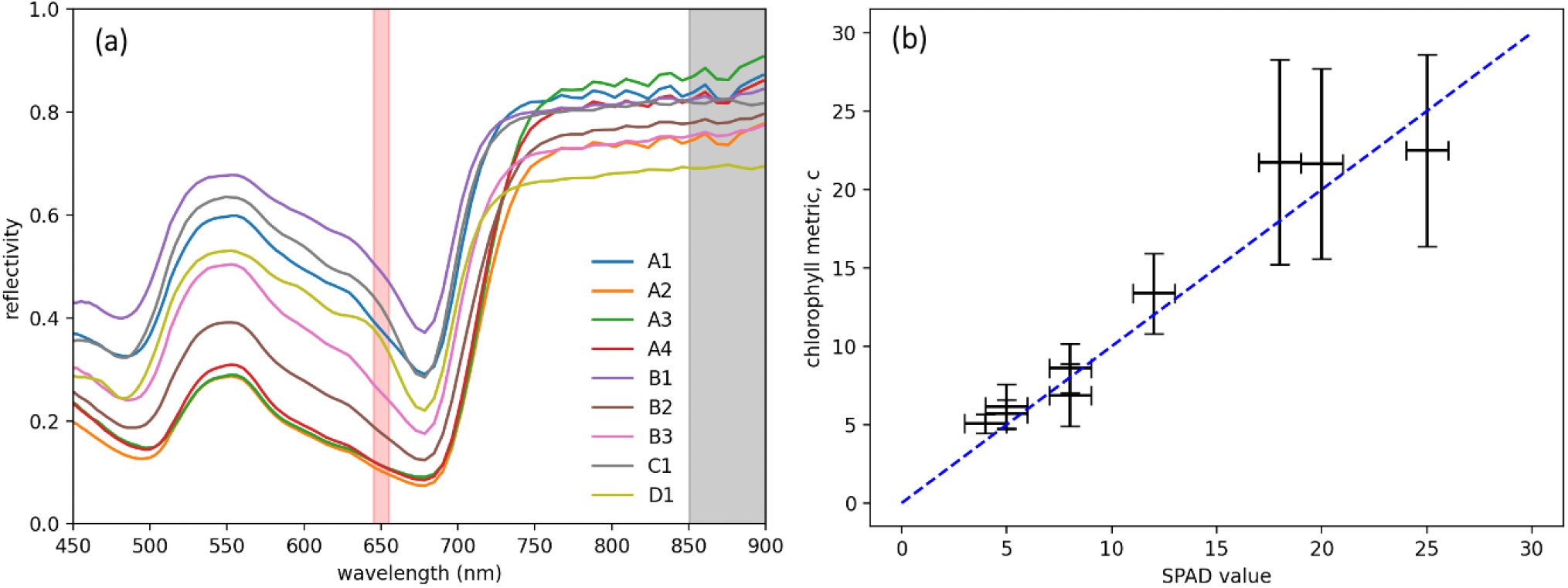
Quantifying chlorophyll content in lettuce leaves using hyperspectral data. (a) Mean spectra extracted from each lettuce leaf ROI. Shaded regions indicate the wavebands using for calculating a ‘SPAD-like’ chlorophyll metric, *c*. (b) Computed chlorophyll metric values plotted against SPAD values obtained using a commercially-available chlorophyll meter. Horizontal error bars indicate the chlorophyll meter’s specified accuracy of *±* 1 SPAD, while vertical bars indicate *±* one standard deviation of the dataset used to calculate the mean. To guide the eye, a blue dashed line, representing *c*=SPAD value, is also plotted.

Following collection of the hyperspectral data, each leaf was also measured using a commercially-available handheld chlorophyll meter (Portable TYS-A Chlorophyll Meter, HSRG), also known as a ‘SPAD meter’. The device contains two LED light sources, one in the red (650 nm) and one in the near-infrared (940 nm). The meter operates by measuring the transmission of light through a 3 x 3 mm region of the leaf at these two wavelengths, and computing the ratio of the two transmitted intensities to return a chlorophyll metric, known as a ‘SPAD’ value.^17^ Using this device, ten measurements were performed within each leaf ROI, and the resulting mean values for each region are presented in Table 2.

**Table 2.** ‘SPAD’ measurements of lettuce leaf regions of interest (ROI), obtained using a handheld chlorophyll meter. Each value is the mean of ten measurements taken within the ROIs indicated in Figure 6.

| Leaf ROI | ‘SPAD’ value |
| --- | --- |
| A1 | 8 |
| A2 | 18 |
| A3 | 25 |
| A4 | 20 |
| B1 | 4 |
| B2 | 12 |
| B3 | 8 |
| C1 | 5 |
| D1 | 5 |

To enable comparison with the SPAD meter, we use Living Optics hyperspectral data to compute a ‘SPAD-like’ chlorophyll metric, *c*, for the lettuce leaf reflectance measurements, as defined below:

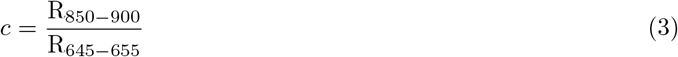

where *R*_850*−*900_ is the mean reflectance value in the range 850 - 900 nm (shaded grey in Figure 7(a)), and *R*_645*−*655_ is the mean reflectance in the 645 - 655 nm range, shaded red in same the figure.

The red band in this metric is selected to centre on 650 nm, in order to mimic the central wavelength probed by the SPAD meter. However, given that the hyperspectral data ranges up to 900 nm, it is not possible to match the SPAD meter’s near-infrared wavelength of 940 nm. Instead, a central wavelength of 875 nm was chosen for the near-infrared band, as for the previous metric (equation 2). Given that the near-infrared response is expected to be largely flat, and noting the fluctuations in the spectral response in this range, a relatively large bandwidth of 50 nm is used, in order to average out the effects of noise in this region.

As shown in Figure 7(b), the proposed chlorophyll metric shows good correspondence with the SPAD readings, within the range of values measured. This indicates the potential of the Living Optics camera to be used as a chlorophyll meter, and in this context, it may offer several advantages over a conventional SPAD meter. Firstly, the reflectance-mode operation allows measurements to be made without physical contact with the plant leaves, and without the need to manually position samples in the device for assessment. As it is an imaging system, rather than a point - spectroscopy device, it is also possible to simultaneously gather information about different parts of a leaf or plant. Furthermore, the video-rate nature of the device could enable continuous, or even autonomous, monitoring of plants *in situ*.

In the next section, we describe early work on an ecological monitoring project that aims to exploit these advantages to enable automated monitoring of grassland and woodland habitats using hyperspectral imaging.

## 4 ECOLOGICAL MONITORING

In this section, we explore the application of the Living Optics camera for monitoring the health and drought resilience of grassland ecosystems. The experiment is located at the Upper Seeds field site (51°46’16.8”N 1°19’59.1”W, 155 m a.s.l) in Wytham woods, Oxfordshire, UK. The site is a calcareous temperate grassland characterised by poorly developed (300 - 500 mm depth) alkaline soils,^37^ with a daily average temperature ranging between -5 °C and 26 °C (2016 - 2020), and daily total precipitation of 0 - 40 mm (2016 - 2020). The results presented here constitute preliminary work to explore a range of analysis methods enabled by the camera, and their suitability for the *in situ* study of plant health. To this end, we collected three test samples of the forb *Potentilla reptans* from Wytham Woods, one from each of three treatment plots described in Table 3. Samples were collected from the site and measured in the laboratory immediately on return, approximately 3 hours later, to secure the measurement of biologically viable tissue. The samples were imaged using a Living Optics hyperspectral camera and a broadband halogen light source (300 - 500 W Fresnel Tungsten Spotlight, HwaMart Ltd, UK). A piece of white material (Tyvek substrate, DuPont, USA) was included in each image to act as a white reference object for reflectance conversion. In the sections below, we discuss a range of computational analysis techniques applied to the hyperspectral data, and demonstrate their use for analysing plant samples.

**Table 3.** Description of treatment plots selected for sample collection at Wytham Woods.

| Treatment plot | Description |
| --- | --- |
| Control | No experimental treatment applied |
| Drought | 50% of precipitation is intercepted by rain shelters, transmitted via gutters and collected in containers situated next to each shelter. |
| Irrigation | The 50% precipitation collected from the drought plot is sprayed via a sprinkler system onto the irrigated treatment plots. This design ensures that the precipitation each experimental treatment receives is proportional to the average natural precipitation across the site. |

### 4.1 Technique 1: Resolution enhancement

In a single snapshot, the hyperspectral arm of the Living Optics camera samples 4,384 spatial points across the scene as illustrated in Figure 8(a). However, the video-rate nature of the camera, together with the high-resolution ‘scene view’ information provided for each frame, enables techniques for generating datasets with far higher spatial resolution.

**Figure 8.**
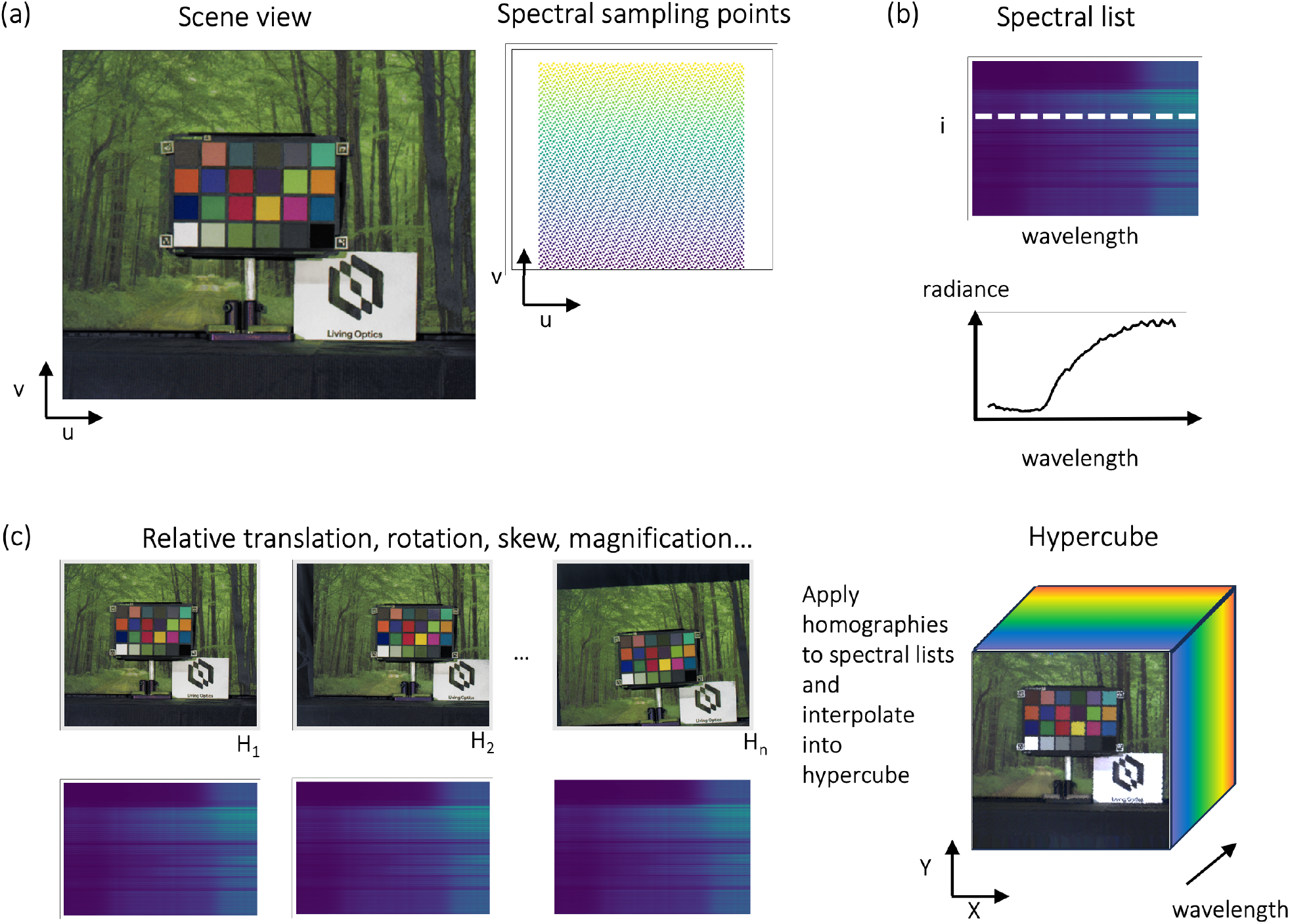
Diagram illustrating how a video acquisition of the Living Optics camera can be used to enhance the spatial resolution of a hyperspectral data cube. (a) Each snapshot gives a scene frame view and a list of spectra shown in (b) originating from specific spectral sampling points that uniformly cover the frame. (c) The homographies of each frame are inferred from the scene view and applied to the lists of spectra to generate a radiance hyperspectral data cube. Note the magnitude of the displacements are exaggerated in (c) for clarity.

This spatial resolution enhancement can be achieved by introducing relative motion between the scene and the camera during video capture, allowing the hyperspectral arm to uniquely sample more points in the global (object space) coordinate frame. Examples of suitable camera motion include panning the camera whilst mounted on a tripod, smooth handheld motion, or natural vibrations of the hand or mounting surface. If the object of interest can be moved, and the scene is relatively flat, it will also be equivalent to move the object. Examples of this include moving a microscope slide on an (x,y) translation stage or moving an object on a conveyor belt.

By algorithmically tracking the movement of key features in the RGB ‘scene view’ image, the magnitude and direction of motion can be inferred. This approach allows the estimation of the global locations of the sampled points in each frame. We note that a similar method of incorporating information from a wide-field sensor has been used to coregister information from a line-scanned hyperspectral endoscopy system for gastrointestinal imaging.^38^ Using standard interpolation techniques, the estimated global coordinates and measured spectra over an entire acquisition can be used to generate a high resolution hyperspectral data cube as described in Figure 8 (b&c).

The method can be described mathematically as follows: two planes can be related geometrically by a projective transformation, or planar homography **H**∈ ℝ^(3×3)^, up to a scaling. The homography matrix encapsulates information about relative shear, translation, rotation and magnification. We first define a global coordinate system (*X, Y*) that describes a static reference plane. We then consider a moving camera imaging that plane in space. If the homographies **H**_**n**_ from the image plane *n* to the reference plane are known, we can apply a projective coordinate transform to map measured spectra into the global coordinate space:

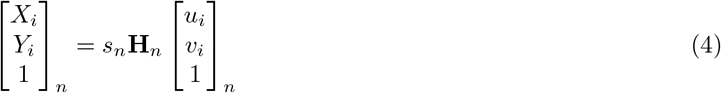

where *s*_*n*_ is a scaling factor and (*X*_*i*_, *Y*_*i*_)_*n*_ and (*u*_*i*_, *v*_*i*_) is the location of the *i*^th^ spectra in global coordinates and the sensor space coordinates respectively.

In our implementation, we define our global coordinate system (*X, Y*) to coincide with the scene in the first frame. The per-frame homography matrices of the *n*^th^ frame relative to the first can then be estimated using readily available computer vision algorithms. The scale-invariant feature transform (SIFT) algorithm is first used to detect features in each of the images.^39^ The detected features are then passed into a Brute-Force matcher, that selects the closest feature descriptor in the pair and matches them. The matched points can then be used to obtain **H**_*n*_, here using an OpenCV method.^40^ Using this information, we then transform all spectra into the global coordinate system. A nearest-neighbours interpolator is used to process the spectra-coordinate pairs, which infills points on a grid by selecting the values of the closest known spectrum. We have chosen to perform interpolation on a square grid, but this approach can be easily extended to generate a hyperspectral data cube of larger dimensions. This may be useful for imaging samples with large aspect ratios. Furthermore, due to the relatively flat nature of our samples, it was appropriate to assume the scene was planar. However, we note that for more complex scenes, it is also possible to create a hyperspectral point-cloud in 3D space using information from both sensors. This method has been applied to hyperspectral mining outcrop^41^ and underground geological mapping,^42^ which enables accurate illumination correction and information fusion with digital twins of the scene.

Figure 9 shows an example output of the resolution enhancement technique, applied to a hyperspectral video of a *Potentilla reptans* (creeping cinquefoil) sample collected from the drought treatment plot at Wytham Woods. Subtle features such as leaf veins, which cannot be resolved in the single snapshot image (b), are clearly resolved in the resolution-enhanced dataset (c). Thus, this technique provides significant advantages for the application of the camera in detailed analyses of plant samples. To further illustrate this advantage, Figure 10 shows NDVI spectral index maps for each of the three *Potentilla reptans* samples, calculated from their resolution-enhanced data cubes. To obtain the relative motion required during datacapture, the camera was loosely mounted on a tripod and then panned and tilted by hand with a magnitude of *<*10 degrees in all directions. The obtained resolution enhancement allows millimetre-scale defects and leaf features to be analysed hyperspectrally.

**Figure 9.**
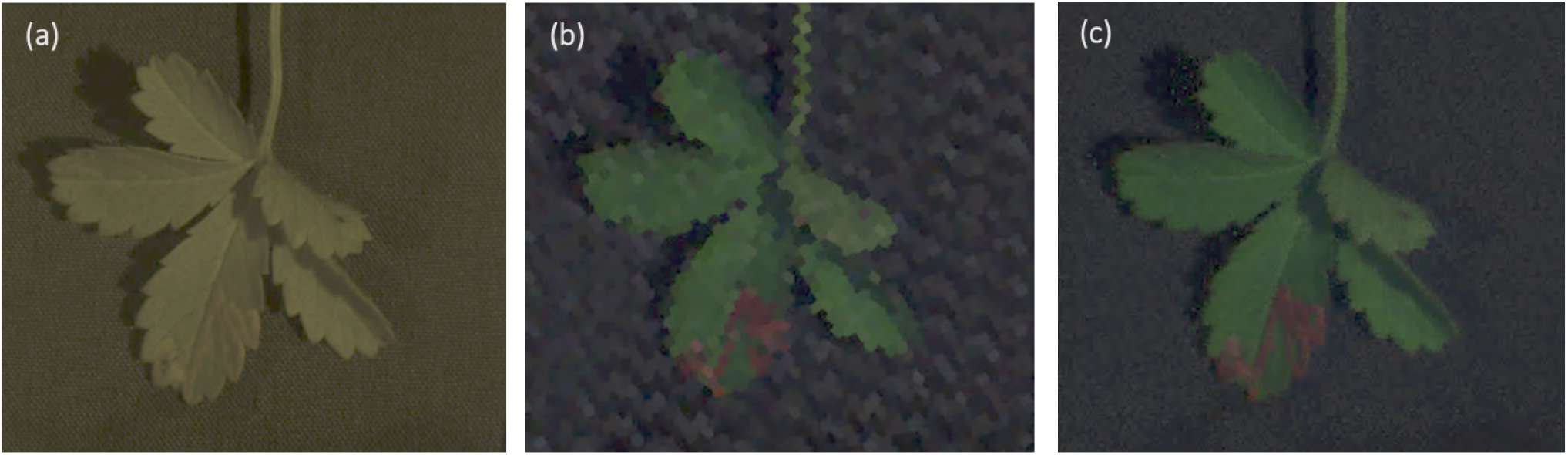
Resolution enhanced data of a *Potentilla reptans* sample collected from the drought treatment plot of the ecological monitoring site at Wytham Woods, UK. (a) ‘Scene view’ RGB image of the sample. (b) An RGB visualisation of the hyperspectral information from a single frame, and (c) the corresponding high-resolution dataset generated in post-processing.

**Figure 10.**
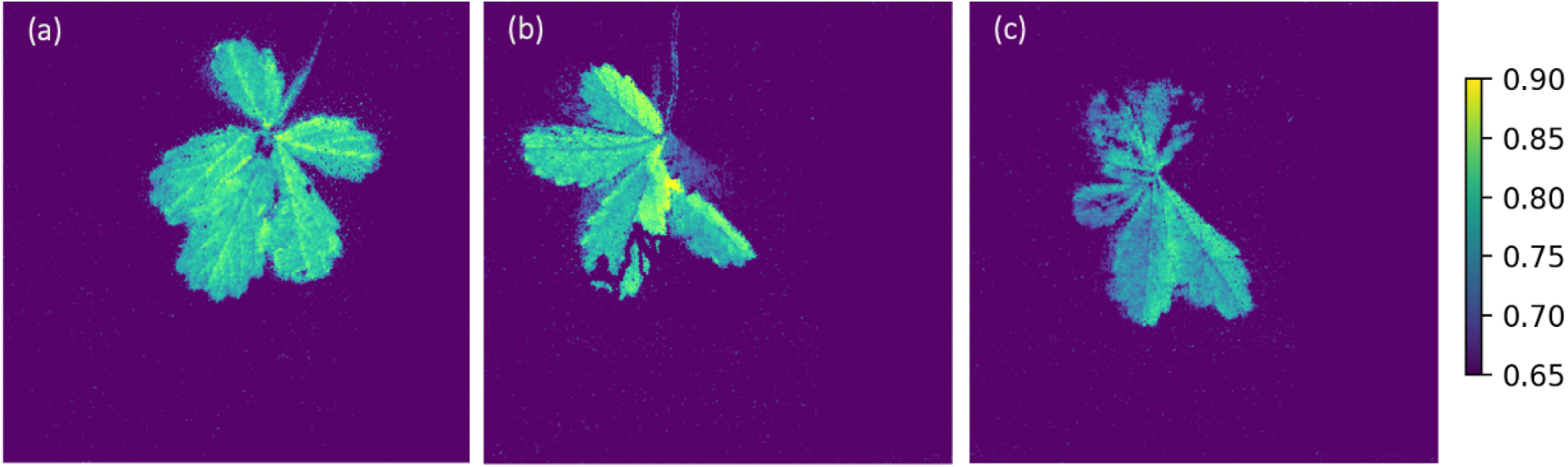
NDVI spectral index maps calculated for plant samples collected from (a) control plot, (b) drought treatment plot, and (c) irrigation treatment plot, constructed using resolution-enhanced datasets. The NDVI calculation was performed using a red waveband of 660-680 nm, and near-infrared band of 790-810 nm. The colourscale limits are optimised to give the best contrast within the leaf regions of the image.

### 4.2 Technique 2: Combining machine learning-based image segmentation with spectral information

The high spatial-resolution ‘scene view’ information can also be used as an input to machine learning and computer vision algorithms, potentially enabling a vast range of advanced analysis techniques to be applied to the hyperspectral data. In the example shown in Figure 11, a widely-used and freely-available deep-learning based zero-shot segmentation algorithm^43^ is applied to the scene view. The algorithm returns binary masks corresponding to different objects in the scene. As a post-processing step, the masks with too few pixels, as well as too many pixels, are removed. Remaining masks indicate the positions of the leaves in the image, and therefore, intersecting the mask locations with their corresponding spectral information allows for automatic extraction of the leaf spectra for each image. However, due to the zero-shot nature of the segmentation method, the result is vulnerable to false positives. This is illustrated in Figure 11(b), where it can be seen that a shadow in the top left of the image has been incorrectly classified as a leaf. In Section 4.3, we discuss alternative segmentation approaches that make use of the spectral information within the data cube. Combining such spectral approaches with RGB-based image classification is expected to increase the accuracy of object classification, and is a matter for future investigation.

**Figure 11.**
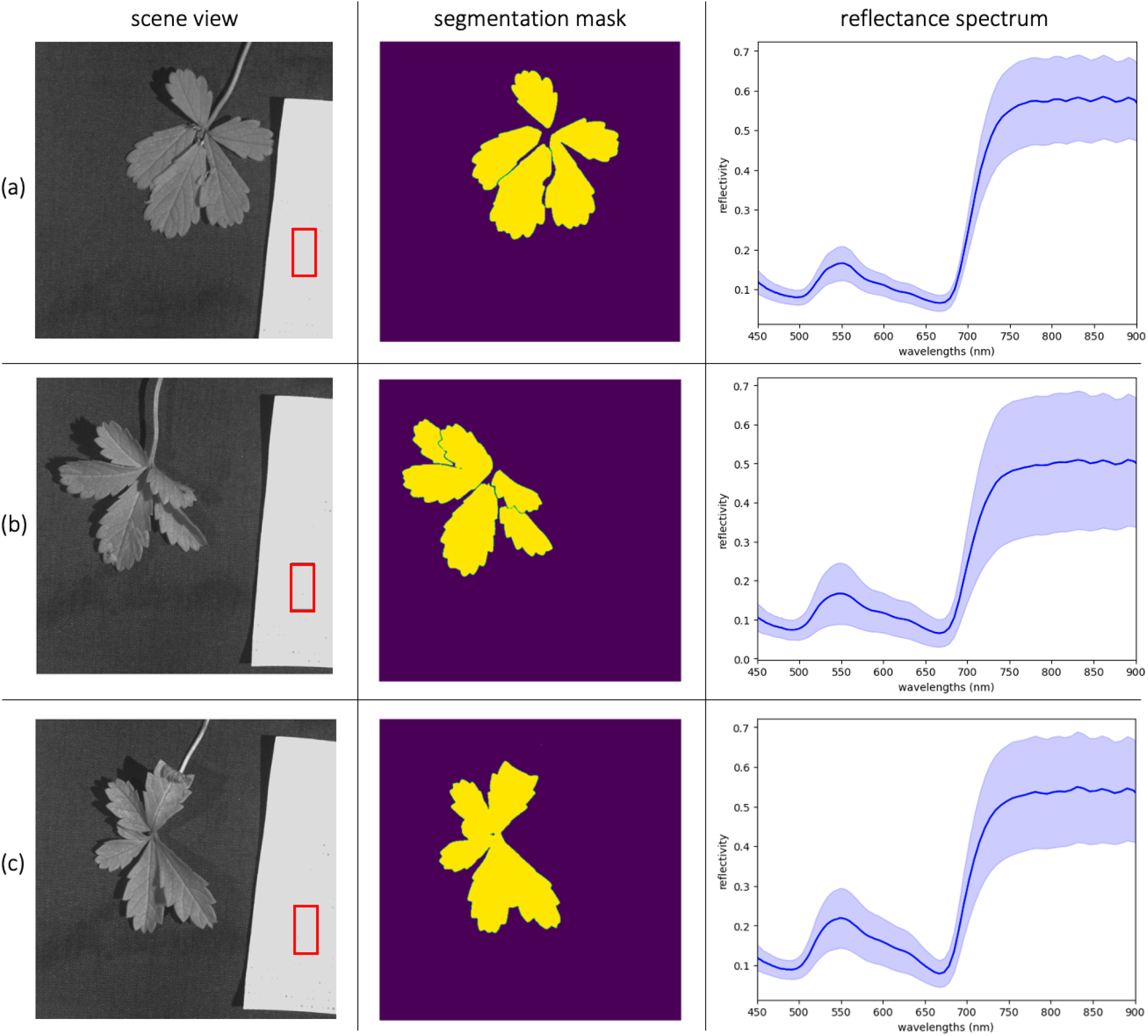
Machine learning image segmentation applied to Living Optics hyperspectral data to extract spectral information from leaf regions. Panel 1: ‘scene view’ RGB images, rendered in greyscale; Panel 2: segmentation masks generated by a deep learning algorithm (Segment Anything, META AI)43 and filtered by area to return only masks relating to leaves. Panel 3: reflectance spectra extracted for the leaf regions. The blue solid lines indicate the mean spectrum calculated from all of the spectra from the extracted region. The shaded region indicates one standard deviation of the data above and below the mean. This analysis is applied to samples from (a) control, (b) drought, and (c) irrigation treatment plots.

Applying the image segmentation analysis described above to the Wytham Woods data, it is observed that all samples show a trough in their reflectance spectrum at approximately 670 nm, consistent with the strong absorption of light by chlorophyll molecules, up to around 700 nm.^11^ All spectra also exhibit a peak at 550 nm, in accordance with the overall green colour of the leaves.^1^ However, subtle differences are seen in the spectral responses of the different plant samples. For example, the sample collected from the irrigation plot shows a broader 550 nm peak, that extends into the yellow region of the spectrum. This pattern is consistent with the appearance of the leaf, which shows regions of yellow/brown discolouration. It is also seen that the 670 nm trough for this sample is not as deep as for the control and drought samples, which may indicate reduced chlorophyll absorption, consistent with the known effects of this treatment at the examined site.^44, 45^

It is also interesting to consider the differences in reflectance in the near-infrared region of the spectrum. High reflectivity in the 700 - 1100 nm range is caused by intercellular light scattering.^1^ Here, the drought and irrigation samples are less reflective than the control sample, which could indicate differences in tissue structure and health, also supported by functional-trait approaches at this site.^44^

Given the small number of samples tested here, it is not possible to draw conclusions as to the cause of the differences between these spectra, or to relate these to their growth environments. However, hyperspectral data cubes collected in this way are rich in information and ripe for further exploration. In the following section, we examine the fine features of the samples to further characterise the leaves’ spectral response.

### 4.3 Technique 3: Spectral angle analysis

In Section 4.2, hyperspectral images were segmented based on shape, through the application of AI computer vision techniques. An alternative approach is to segment the images based on regions that are spectrally alike.

In hyperspectral imaging analysis, a popular metric of spectral similarity is the ‘spectral angle’, in which two spectra are treated as if vectors, and the angle between them is computed.

Mathematically, the spectral angle between a ‘signal’ spectrum **t** and a ‘reference’ spectrum **r** is given by:

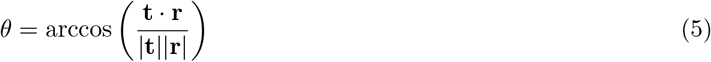

or equivalently:

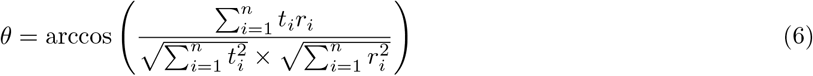

where *n* is the number of wavelength channels in the spectral data.^46^

In this work, spectral angle maps were computed to distinguish green regions from discoloured regions of the leaves. For each sample, two spectral angle maps were created, one calculated relative to a point within the green leaf region, and one relative to a point inside the discoloured region. By applying a threshold to these maps, a binary mask could be produced, and spectra from the two classes (green vs. discoloured) could then be extracted and analysed.

Figure 12(a) shows the results of this analysis for the drought plot sample. It can be seen that the mean spectrum extracted from the red/brown discoloured regions is quite distinct from that of the green leaf regions. Firstly, the discoloured region peak is located in the red region of the spectrum, rather than in the green. This is in accordance with its red/brown colouration, as seen in the bottom right inset. Notably, for the discoloured region, the trough associated with chlorophyll absorption is less pronounced, and the near-infrared reflectivity is lower. As previously discussed, this may suggest lower chlorophyll content in the discoloured region, and possible differences in intercellular structure.

**Figure 12.**
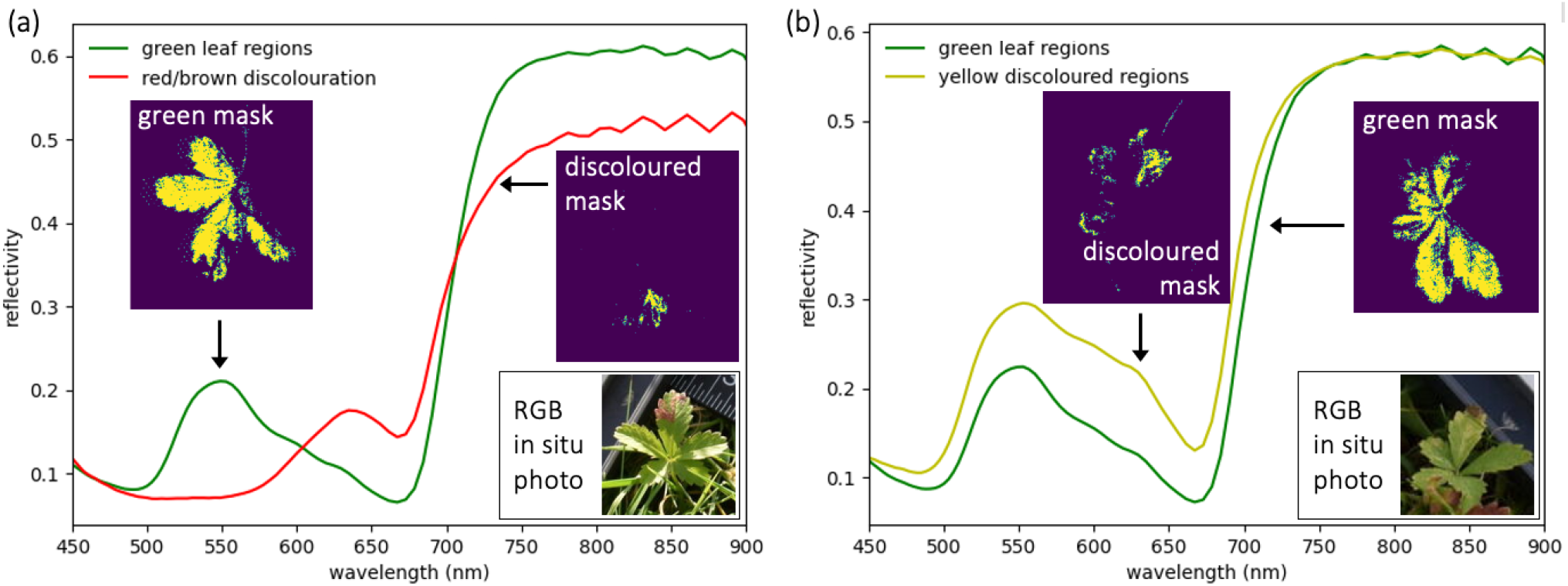
Spectral angle analysis for (a) drought plot sample; and (b) irrigation plot sample. The mean reflectivity spectrum for each extracted region is plotted in the main panel, and the thresholded spectral angle masks (applied for spectra extraction) are shown as insets. A photograph of the corresponding plant, growing *in situ* prior to collection, can be seen in the bottom right inset of each panel.

The results of a similar analysis for the irrigation sample are shown in Figure 12(b). In this case, there is little difference in near-infrared reflectivity for the green and yellow/brown discoloured regions. However, the 550 nm peak is higher and broader, extending into the yellow region, and the chlorophyll absorption trough is less pronounced than for the green leaf regions.

These preliminary results demonstrate the interesting and valuable information that can be obtained using imaging spectroscopy for plant analysis. In particular, the ability to spectrally characterise fine-scale leaf features, as illustrated in this section, opens up possibilities for the use of the Living Optics camera for automated detection and monitoring of plant stress and disease in both agricultural and ecological contexts. In the next section, we discuss future work to explore such applications and to support scientific discovery in these fields.

### 4.4 Future work

Our novel technology allows for fast, cheap, portable, high-resolution quantification of hyperspectral information in natural as well as laboratory conditions. Monitoring natural systems is the basis for a better understanding of the ecological relationships that sustain biodiversity and ecosystem services. However, doing so via classical approaches can prove challenging on multiple axes: price, time, validation, etc. The next frontier in Ecology involves the creation of ‘Digital Twins’,^47^ fully computerised models of natural systems where flows of information are dynamically tracked, and where simulations (e.g. droughts) can safely be conducted to understand key properties of the ecosystem, such as productivity, resilience, and viability. Coupling our hyperspectral camera with existing motion-based platforms such as autonomous robots, drones, and low-elevation plane flights, may unlock the opportunity to deliver a fully functional, dynamic, 3D model of even complex ecosystems, such as tropical forests. Furthermore, the application of hyperspectral image analyses for the monitoring of ecosystems is not limited to the Plant Kingdom. Indeed spectral image analyses is starting to be adopted for the monitoring of animals.^48, 49^

## 5 CONCLUSION AND OUTLOOK

In this paper, we have presented a newly-available, video-rate, snapshot hyperspectral camera, and explored the utility of this system for applications relating to plant health. Promising results have been demonstrated for the quantification of leaf chlorophyll content via a ‘SPAD-like’ spectral metric, as well as the display and analysis of spectral indices relating to plant health, such as NDVI and MCARI. The camera provides spectral information across the 400 - 900 nm range, alongside high-resolution RGB imagery and powerful edge compute, enabling live spectral analysis. A range of analysis techniques that leverage this combination of features has also been presented in this work, including homography-based resolution enhancement, the application of deep-learning image segmentation, and spectral angle analysis. Preliminary investigations applying these techniques to analyse plant samples, collected as part of an ecological study, suggest that these approaches can provide a rich source of valuable information that may be useful for monitoring natural ecosystems. In future work, we look to explore the integration of the camera with drones and quadruped robots, with the aim of enabling autonomous data collection and supporting the creation of a Digital Twin of the ecological monitoring site at Wytham Woods, UK.

## ACKNOWLEDGMENTS

MQ, EF, and RSG were supported by an EPSRC grant and a NERC Pushing the Frontiers grant to RSG.

## Notes

### Competing Interest Statement

The authors have declared no competing interest.

### Summary of Updates

Correction to equation (3)

